# Architecture of the chromatin remodeler RSC and insights into its nucleosome engagement

**DOI:** 10.1101/804534

**Authors:** Avinash B. Patel, Camille M. Moore, Basil J. Greber, Jie Luo, Jeff Ranish, Eva Nogales

**Author notes:** Contributed equally (Avinash B. Patel, Camille M. Moore, Basil J. Greber).

## Abstract

Eukaryotic DNA is packaged into nucleosome arrays, which are repositioned by chromatin remodeling complexes to control DNA accessibility^1,2^. The *Saccharomyces cerevisiae* RSC (<u>R</u>emodeling the <u>S</u>tructure of <u>C</u>hromatin) complex, a member of the SWI/SNF chromatin remodeler family, plays critical roles in genome maintenance, transcription, and DNA repair^2–4^. Here we report cryo-electron microscopy (cryo-EM) and crosslinking mass spectrometry (CLMS) studies of yeast RSC complex and show that RSC is composed of a rigid tripartite core and two flexible lobes. The core structure is scaffolded by an asymmetric Rsc8 dimer and built with the evolutionarily conserved subunits Sfh1, Rsc6, Rsc9 and Sth1. The flexible ATPase lobe, composed of helicase subunit Sth1, Arp7, Arp9 and Rtt102, is anchored through the interactions between the N-terminus of Sth1 and the core. Our cryo-EM analysis also shows that in addition to the expected nucleosome-Sth1 interactions, RSC engages histones and nucleosomal DNA through one arm of the core structure, composed of Rsc8 SWRIM domains, Sfh1 and Npl6. Our findings provide structural insights into the conserved assembly process for all members of the SWI/SNF family of remodelers, and illustrate how RSC selects, engages, and remodels nucleosomes.

## Main

Eukaryotes have four major families of chromatin remodelers: SWI/SNF, ISWI, CHD, and INO80^2^. Each of these remodelers plays distinct roles based on how they select and affect target nucleosomes. Together, these remodelers give rise to the distinct chromatin landscapes observed in eukaryotic cells and determine how genetic information is organized, replicated, transcribed, and repaired^5^. In *S. cerevisiae* there are two members of the SWI/SNF family of chromatin remodelers: RSC and SWI/SNF^3,4^. RSC is essential for viability and is ten times more abundant than SWI/SNF^4^. While both complexes play a role in remodeling nucleosomes during transcription initiation, RSC is also involved in transcription-independent processes such as mitotic division, double stranded break repair, and telomere maintenance^6–12^.

RSC is a ∼1-MDa complex composed of 17 proteins, with two copies of Rsc8 and one copy of either Rsc1 or Rsc2 (Fig. S1)^4,13^. In order to determine the structure of RSC, we purified the complex from *S. cerevisiae* and performed cryo-EM analysis (Fig. S2). We find that RSC is composed of five main lobes, three that form a rigid core (head, body and arm) and two that are flexibly attached (leg and tail) (Fig. 1a, b; Fig. S3, S4). We determined the structure of the core to ∼3.1 Å and mapped 14 proteins within this region: Rsc1/2, Rsc3, Rsc4, Rsc6, Rsc8 (2 copies), Rsc9, Rsc30, Rsc58, Ldb7, Npl6, Htl1, Sfh1, and Sth1 (Fig. 1c, d, e). Our negative stain analysis of the yeast SWI/SNF complex shows that, like RSC, it has head, body, and arm regions that define a rigid core, and a flexible leg that occupies similar overall positions (Fig. S5). However, SWI/SNF lacks a tail lobe, indicating that while RSC and SWI/SNF share a conserved core architecture, RSC features additional regulatory domains (Fig. S5).

**Figure 1.**
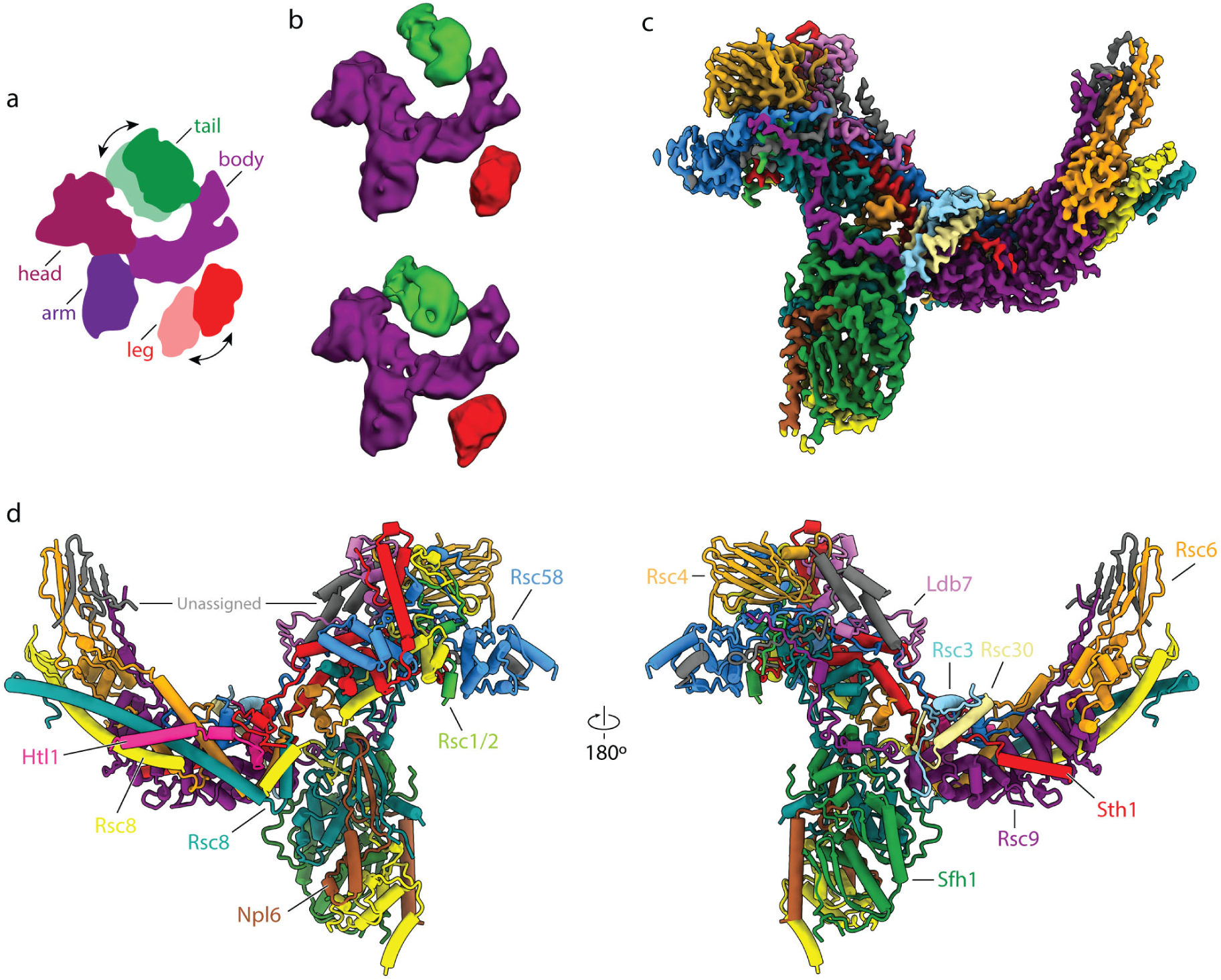
Structure of the RSC core. **a**, Cartoon representation of RSC showing its five main lobes. The head, body and arm lobes form the core of the complex, while the tail and leg lobes are flexible. **b**, Cryo-EM reconstructions of RSC with the tail (green) and leg (red) lobe in two different conformations with respect to the core (purple). **c**, Cryo-EM reconstruction of the RSC core with individually subunits colored. **d**, Model of RSC shown from the front (left) and back (right) views.

To further confirm our model, we used bis(sulfosuccinimidyl)suberate (BS3) to chemically crosslinked the RSC complex and performed mass spectrometry analysis (Fig. S6). We identified 780 unique interlinks between different subunits and 617 unique intralinks within the same subunits. About 90% (151/168) of mappable crosslinks in our model of the RSC core structure are within 38 Å distance (Fig 1a, b, Fig. S6b).

The RSC core structure is critically defined by Rsc8, Rsc6, Rsc9, Rsc58, Sth1, and Sfh1 (Fig. 1d, e). Five of these proteins (Rsc8, Rsc6, Rsc9, Sth1, and Sfh1) are evolutionarily conserved throughout the eukaryotic SWI/SNF family and comprise a considerable amount of the RSC core density (Fig. 2a)^14,15^. In particular, an asymmetric Rsc8 dimer defines the backbone for the complex, scaffolding the three core lobes and contacting all core proteins except Rsc3 and Rsc30 (Fig. 2b). Previous work has shown that Rsc8 homologs are critical for the integrity of the human SWI/SNF family complexes, supporting the idea that this protein is the structural backbone for the entire family of remodelers^16^.

**Fig. 2.**
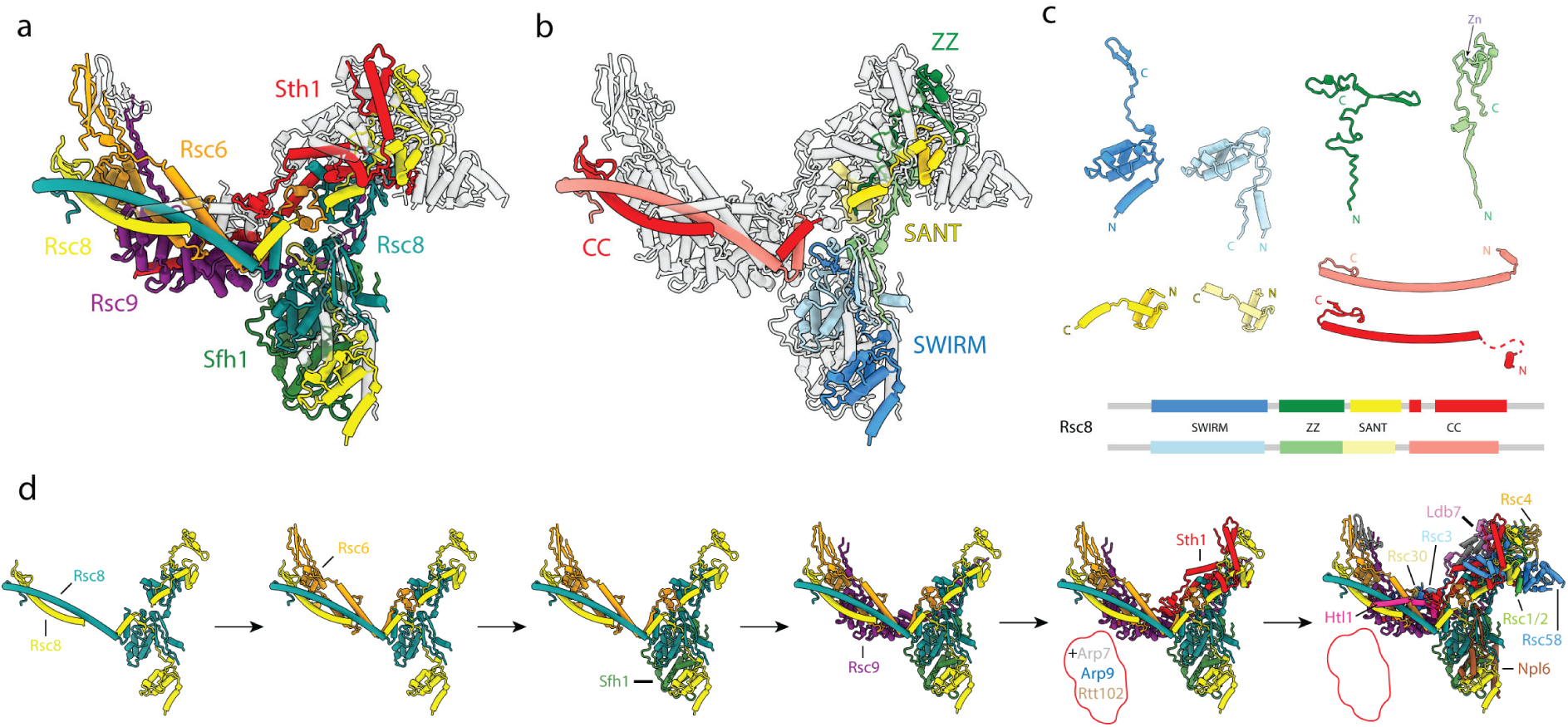
Scaffolding of the RSC core. **a**, Core of RSC with the evolutionarily conserved subunits colored. Non-conserved subunits are shown in transparent grey. Subunit labels are shown with colors corresponding to those in the structure. **b**, Structure of the core with only the Rsc8 dimer colored according to its four domains, blue to red from N to C terminus. **c**, Individual domains of the two copies of Rsc8, aligned and tiled, with domain map shown below. **d**, Proposed assembly process of RSC highlighting the initial stages, which involve only evolutionarily conserved subunits.

Rsc8 has four conserved domains: SWIRM, ZZ, SANT, and coiled coil (CC) (Fig. 2c). The Rsc8 N-terminal SWIRM domains interact with Sfh1 to define the arm. Although the Rsc8 SWIRM domain has been reported to bind DNA, the proposed binding site mediates protein-protein interactions within RSC, thus making it unlikely that this domain is involved in DNA binding^17^. The two ZZ domains assume different folds within RSC: one copy forms two long β-strands and constitutes most of the head density, while the other adopts a zinc-binding ZZ-type fold that connects the arm and head lobes (Fig. 2c). The Rsc8 SANT domains scaffold the head lobe of the core. Notably, the RSC body is built around the two Rsc8 C-terminal CC domains, which are the main site of direct interaction between the two Rsc8 copies.

The pattern of protein-protein interactions observed in our structure provides insight into RSC assembly (Fig. 2d). We propose that the dimerization of the Rsc8 CC domains is the first step in RSC complex assembly, which is likely followed by the binding of Rsc6 to this CC dimer. A tandem strand-strand-helix-helix motif within Sfh1 would then bring the two SWIRM domains of Rsc8 together. Subsequent assembly steps likely involve binding of Rsc9 and Sth1. The armadillo repeat domain of Rsc9 binds the Rsc8 CC and Rsc6 to form most of the body lobe, while the scaffolding domains (I and II) of Sth1 contribute to both the head and body regions and exit towards the flexible leg lobe. Immediately following the scaffolding domains of Sth1 is the HSA helix, which is known to interact with Arp7 and Arp9. These subunits, along with Rtt102, form the Arp module that we propose constitutes most of the leg (see below)^18,19^. It is important to note that this initial assembly process only involves evolutionarily conserved subunits and thus is likely conserved for all SWI/SNF chromatin remodelers (Fig. S5a)^15,16^. This conclusion is further supported by the structural similarities we observe between the yeast SWI-SNF and RSC complexes, and by the previous mass spectrometry analysis of the assembly of human BAF complexes^16^.

The assembly of RSC likely continues with the addition of the yeast specific factor Npl6, which completes the arm lobe, and RSC specific subunits Htl1, Rsc58, Rsc1/2, Rsc4, and Ldb7. These subunits either contribute to the scaffold of the complex (Htl1 and Ldb7), serve to anchor bromodomains to the core (Rsc1/2 and Rsc4) or both (Rsc58). The last two subunits to be recruited to the complex are likely the RSC-specific Rsc3 and Rsc30. These two proteins, which are expected to be globular and helical in fold, appear to be absent in 75% of the RSC particles subset used for the high-resolution refinement (Fig. S1, S7). This agrees with previous findings that showed that there are RSC complexes that lack Rsc3 and Rsc30^20^. During our image analysis, we found that the presence or absence of Rsc3 and Rsc30 correlates well with the presence or absence of most of the tail lobe, making the Rsc3 and Rsc30 subunits the most likely constituents of this very flexible lobe (Fig. S7a). The fact that there are no equivalent subunits in the yeast SWI-SNF complex, for which the tail domain is missing, further supports this assignment (Fig. S5a).

In order to shed light on the interaction of RSC with its substrate, we obtained an 19Å resolution map of RSC bound to a nucleosome core particle (NCP) modified with H3K4me3 and H3K(9/14/18)ac (Fig. 3a,b; Fig. S8). We were able to accurately fit our structure of the RSC core within this map, along with models of the Arp module (Arp7, Arp9, Rtt102 and Sth1-HSA) and the NCP (Fig. 3c) ^18,21^. About 88% (206/233) of mappable crosslinks in our model of the RSC-Arp module structure are within 38 Å distance (Fig 3b, Fig. S6c). We do not observe any density for the ATPase domain of Sth1; however, based on the structure of the Sth1 homolog Snf2 bound to the NCP, we expect the catalytic subunit to bind super-helical location-2 (SHL2) (Fig. 3c)^21^. In the resulting model, the N and C termini of adjacent Sth1 domains are at the right distance to connect through the short linkers between them (Fig. 3c). Our map shows four regions of interaction between RSC and the NCP. Two involve the arm of RSC and include the C-terminal portion of the Snf5 domain of Sfh1 and part of Npl6 (Fig. 3d). Within the RSC-NCP map, the Snf5 domain appears to continue towards the NCP to interact with the acidic patch (Fig. 3c; Fig. RSC-NCP2). This is supported by the fact that the Snf5 domain has a highly conserved set of basic residues at its C-terminus and deletion of this subunit results in less efficient chromatin remodeling activity of SWI/SNF complexes (Fig. 3c)^22^. The second point of NCP contact occurs between the arm of RSC and SHL4 (Fig. 3c). Based on the position of this interaction, we predict that the unmodeled parts of Npl6 form this interaction. The third RSC-NCP contact point involves the Arp module in the leg. This connection appears to be between Rtt102 and the H2A tail (Fig. 3c). Similar contacts to the three just described have been shown to occur also in the SWI/SNF complex, involving the homologous proteins for Sfh1 (Snf5), Npl6 (Swp82), and Rtt102^23^. The fourth and last point of contact between RSC and the NCP appears to be more dynamic and involves the tail lobe and either the H3 tail or the extranucleosomal DNA (Fig. 3c). We observe two distinct modes of NCP binding involving this region, one where the whole tail is swiveled towards the nucleosome, and one where only a small region emanating from the tail reaches towards the NCP (Fig. 3a, b, d). Altogether, the core of RSC appears to bind the nucleosome in a well-defined orientation that places SHL2 in position to interact with the catalytic domain of Sth1. The fact that we observe nucleosome-bound RSC, even without the binding of the catalytic domain of Sth1 to the NCP, indicates that the contacts through the RSC core are sufficient for nucleosome engagement. The contacts made by the RSC core with the NCP are likely to contribute to the fidelity of the nucleosome remodeling function of the catalytic domain of Sth1 (see below).

**Fig. 3.**
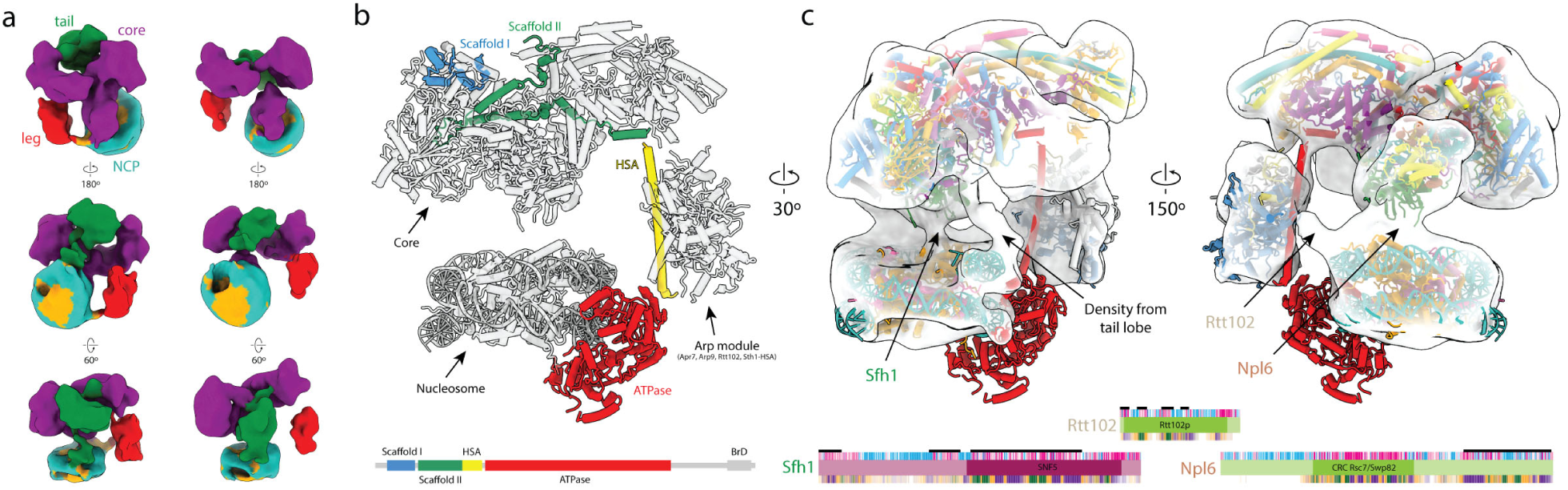
Nucleosome binding by RSC. **a**, Cryo-EM reconstructions of RSC-NCP with the tail (green) and leg (red) and core (purple) contacting the NCP (DNA in teal and histones in orange). In the structure on the left, only a small region of the tail is contacting the NCP, while in that on the right, the entire tail is swiveled towards the NCP. **b**, Model of RSC-nucleosome with domains of Sth1 highlighted blue to red from N to C terminus. Scaffolding domains I and II interact with the core region of RSC, while the HSA helix interacts with the Arp module to form the leg lobe. The ATPase (not visible in our density) is modeled according to the structure of nucleosome-bound Snf2 (ref. ^21^). **c**, Interactions between the RSC core and the nucleosome. Contacts are observed between (i) the acidic patch of the nucleosome and Sfh1, (ii) the nucleosome SHL0 and the tail lobe, (iii) nucleosome SHL4 and Npl6, and (iv) the H2A tail and Rtt102. Domain maps of Sfh1, Rtt102 and Npl6 are shown below. From top to bottom, black bar indicates the regions of the protein that are modeled in our structure, pink-white-blue bar represents sequence conservation according to ConSurf scores (most to least), PFAM predicted domains (shown in bold with labels), and predicted secondary structure (purple for helix, green for stand, orange for coil and white for disordered)^28–34^.

A notable aspect of RSC is that it has six bromodomains, two on Rsc4, two on Rsc1/2, one on Sth1, and one on Rsc58 (Fig. 4a; Fig. S1) that associate with acetylated lysines on histone tails, particularly H3^24^. For comparison, the yeast SWI/SNF and its human homolog BAF have just one bromodomain – on the ATPase subunit – while human PBAF has eight, indicating that these domains may contribute to functional specificity of different classes within the SWI/SNF family (Fig. S5). We could model the bromodomain on Rsc58 as a part of the core RSC region, but the five remaining domains extend from flexible linkers and are not visible in our structure (Fig. 4a). Based on the position of the bromodomain-containing subunits in our RSC-nucleosome model, all of these domains could reach the H3 tails of the engaged nucleosome. Alternatively, because we only observe a single mode of RSC-nucleosome engagement, it is possible that the bromodomains of one RSC complex could interact with different, adjacent nucleosomes. Microarray data has shown that different bromodomains within RSC can recognize different acetylation sites, which could allow the complex to be targeted to a wide variety of genomic loci^25,26^.

**Fig. 4.**
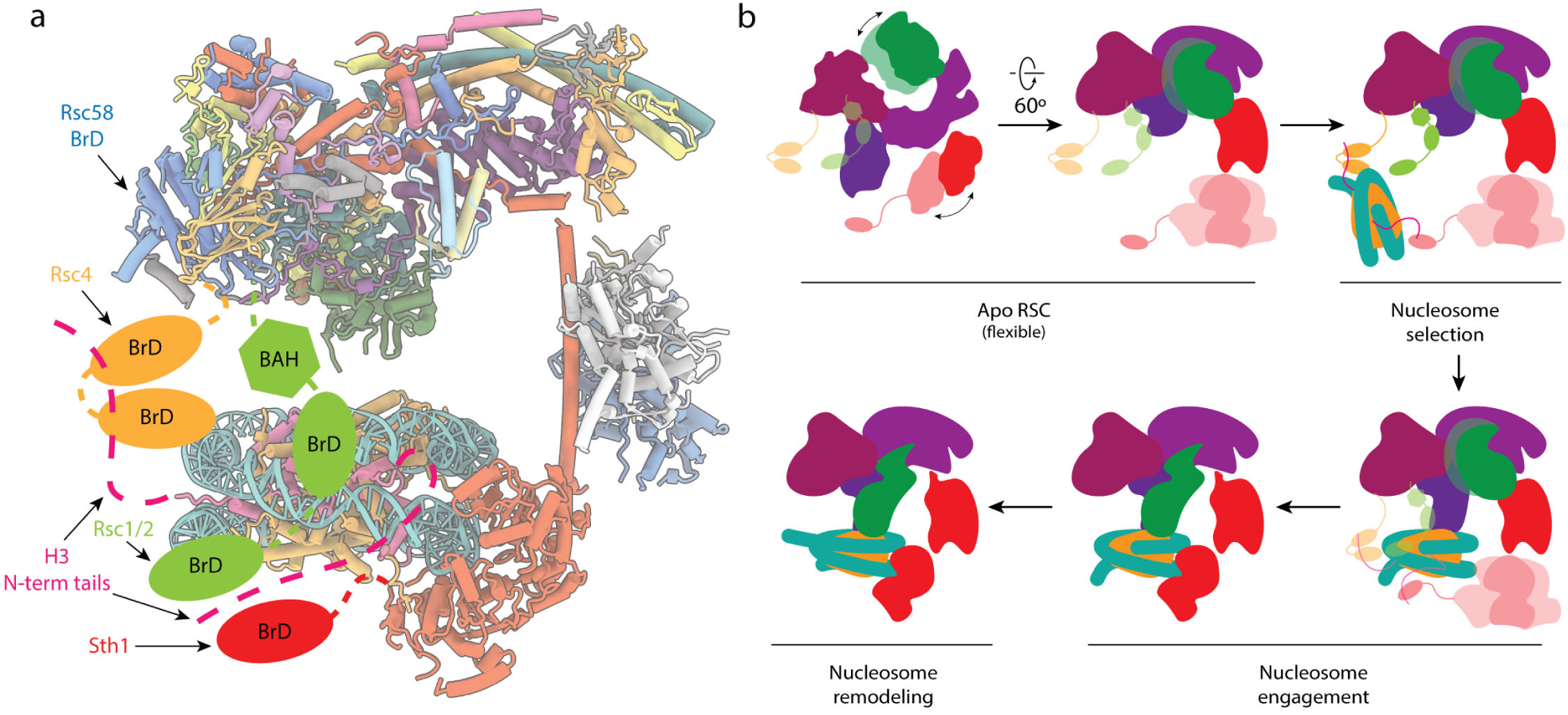
Chromatin interacting domains of RSC and model of nucleosome engagement. **a**, Of the six bromodomains (BrD) present in RSC, only the one in Rsc58 (shown in blue) is rigidly attached to the core and modeled in our structure. The other five (2 on Rsc4, 2 on Rsc1/2, and 1 on Sth1) are flexibly attached to the core of RSC and positioned such that they can interact with histone H3 tails (pink) in the engaged nucleosome. **b**, Mechanistic model of RSC engaging a nucleosome. The apo RSC is shown with its moving tail and leg lobes, and its flexibly attached BrDs. The BrDs bind and select target nucleosomes with acetylated tails. The selected nucleosome engages RSC, first through the arm lobe of the core, which then positions the ATPase domain at the end of the leg so as to be able to bind SHL2. During nucleosome remodeling, the ATPase translocates the DNA while the RSC core holds onto the histone core.

Based on our structure and existing biochemical data, we propose a 4-step mechanism of how RSC functions on chromatin (Fig. 4b). Initial recruitment of RSC to a nucleosome likely occurs through interaction with H3 acetylation marks, since RSC has an increased affinity for these nucleosomes^27^. RSC will then engage the nucleosome through its arm region and place the SHL2 site of the nucleosome in position to bind the flexibly tethered catalytic domain of Sth1. The ATPase domain is then able to translocate the DNA around the histone octamer, while the core of RSC holds onto the histone core through its direct interactions with histones. This process is very likely to apply to all SWI/SNF chromatin remodelers, as they share the same architecture and contain the conserved subunits/domains involved in the steps just proposed. Specifically, all complexes have bromodomains to bind acetylated histones (minimally in the C-terminus of their catalytic subunit), a Sfh1 homolog to bind the acidic patch of the nucleosome, and a catalytic DNA translocase domain to remodel nucleosomes (Fig. S5)^2^.

In summary, our studies provide the detailed structure for the conserved core of SWI/SNF complexes and lead to a model of assembly for this family of remodelers that agrees with previous biochemical data on mammalian complexes. Our work also provides a model of how RSC stably contacts the core histones in a nucleosome via its body module, while engaging nucleosomal DNA via the Sth1 ATPase. Both the structural integrity of the core and the interaction with nucleosomes rely on evolutionarily conserved subunits, while the non-conserved proteins in RSC are likely to play a role in recruitment and regulation of the complex. Future studies will be needed to understand the mechanisms by which individual members of the SWI/SNF family of chromatin remodelers are brought to different genome loci and are distinctly regulated.

## Acknowledgements

We thank A. Iavarone for performing in-gel mass spectrometry data collection and analysis, P. Grob, and D. Toso for electron microscopy support, A. Chintangal and P. Tobias for computing support, and C. Yoshioka and the OSHU Cryo-EM Facility for help with data collection. A portion of this research was supported by NIH grant U24GM129547 and performed at the PNCC at OHSU and accessed through EMSL (grid.436923.9), a DOE Office of Science User Facility sponsored by the Office of Biological and Environmental Research. This work was funded through NIGMS grants R01-GM63072 and R35-GM127018 to E.N., and R01-GM110064 and R01-HL133678 to J.R. E.N. is a Howard Hughes Medical Institute Investigator.

## Author contributions

A.B.P. and C.M.M prepared, collected and processed EM data. A.B.P. and B.J.G built atomic model. B.J.G refined the atomic coordinate model. J.L. and J.R. performed cross-linking mass spectrometry analysis. A.B.P, C.M.M and E.N. wrote paper with input from all authors.

## Competing interests

The authors declare no competing interest.

## Methods

### Protein Purification

SWI/SNF was purified from *Saccharomyces cerivisiae* using a modified TAP purification as described in Nagai et al^35,36^. A strain modified with a TAP tag on the C terminus of SNF2 was obtained from GE Dharmacon and grown at 30 °C in YPD. 20 L of cells were harvested at OD 6 and lysed using a cryo-mill. The ground cells were resuspended in a lysis buffer (50mM HEPES pH 7.9, 25 mM ammonium sulfate, 0.5 mM EDTA, 100 μM zinc sulfate, 5% glycerol, 5 mM DTT, 10 μM leupeptin, protease inhibitor, .01% NP-40) and dounced to ensure homogenization. The lysate was spun at 11,000 g for 20 min. The supernatant was removed and brought to 200 mM ammonium sulfate, followed by an addition of polyethyleneimine (0.2% final concentration) to precipitate DNA. The sample was spun again at 17,000 g for 40 min. Supernatant was removed and brought up to 2.2 M ammonium sulfate, then spun again at 17,000 g for 40 minutes. The protein pellet was resuspended in buffer (lysis buffer with 0 M ammonium sulfate and 2 mM DTT) and the ammonium sulfate concentration was brought up to 400 mM. IgG resin (1 ml packed) was equilibrated and incubated for 4 hours. Resin was washed with 500 mM ammonium sulfate buffer, then 1x with 100 mM ammonium sulfate buffer. Protein was released via TEV cleavage by overnight incubation with 20 μg TEV (MacroLab) in AS-100. Following cleavage, samples were spun to collect supernatant. The CaCl_2_ concentration of the supernatant was brought up to 2 mM and loaded onto a 100 μl equilibrated CBP resin. After 4 hours of incubation, supernatant was removed and the resin was washed 4x (20mM HEPES pH 7.9, 2 mM MgCl_2,_ 10% glycerol, 250 mM KCl, 50 μM ZnCl_2_, 2mM CaCl_2_, 10 μM leupeptin, 1 mM TCEP, 0.01% NP-40). The sample was eluted with one volume elution buffer (20mM HEPES pH 7.9, 2 mM MgCl_2_, 10% glycerol, 250 mM KCl, 50 μM ZnCl_2_, 2 mM EGTA, 10 μM leupeptin, 1 mM TCEP, 0.01% NP-40) for 30 min, then 15 min for each subsequent elution. Samples were aliquoted, frozen in liquid nitrogen and stored at -80 °C.

RSC was purified from *Saccharomyces cerivisiae* using a TAP-tag method as described^35,37^. A strain modified with a TAP tag at the C terminus of STH1 was obtained from GE Dharmacon and grown at 30 °C in YPD. 10 L of cells were harvested at OD 7 and lysed using a cryo-mill. The ground cells were resuspended in a lysis buffer (50 mM HEPES pH 7.9, 250 mM KCl, 0.5 mM EDTA, 100 μM ZnSO_4_, 10% glycerol, 2 mM DTT, 10 μM leupeptin, protease inhibitor, 0.05% NP-40) and dounced to ensure homogenization, followed by an addition of 18 μL of benzonase (Sigma) while spinning on ice. After 10 min, another 18 μL of benzonase were added. Heparin (Sigma) was then added slowly to a final concentration of 0.5 mg/mL and incubated for 10 min. The lysate was then spun for 90 min at 17,000 g. Supernatant was collected and clarified through a column frit, then bound to 2.5 mL packed IgG resin and incubated for 4 hours at 4 °C. After incubation, the supernatant was removed. TCEP was added to the lysis buffer for a final concentration of 0.5 mM and washed 5x over the resin. The sample was released via overnight incubation with 25 μg TEV protease in one volume buffer. Following cleavage, samples were spun to collect supernatant. The CaCl_2_ concentration of the supernatant was brought up to 2 mM and loaded onto 100 μL CBP resin equilibrated five times with wash buffer (20mM HEPES pH 7.9, 10% glycerol, 150 mM KCl, 50 μM ZnCl_2_, 2 mM CaCl_2_, 10 μM leupeptin, 1 mM TCEP, 0.01% NP-40). After 4 hours of incubation, the supernatant was removed and the resin was washed 2x with wash buffer with 750 mM KCl, then 3x with wash buffer with 250mM KCl buffer, then 1x with wash buffer without CaCl_2_. The sample was eluted with one volume (20mM HEPES pH 7.9, 2 mM MgCl_2_, 10% glycerol, 150 mM KCl, 50 μM ZnCl, 2 mM EGTA, 10 μM leupeptin, 1 mM TCEP, 0.01% NP-40) for 30 minutes, then 15 minutes for each subsequent elution. Samples were aliquoted, frozen in liquid nitrogen and stored at -80 °C.

### Chemical Crosslinking Mass Spectrometry

We used the ∼250 ug RSC2 TAP-tag purified RSC complex and crosslinked by 4mM bis(sulfosuccinimidyl)suberate (BS3) at RT for 2 hours. Sample processing and mass spectrometry data analyses were done as described before^16^. After plink2 and Nexus database searches against the RSC subunit sequences, about 6.6% of interlinked spectra and 3.1% intralinked spectra were removed after manually spectrum checking. All the crosslinked spectra can be viewed at https://www.yeastrc.org/proxl_public/viewProject.do?project_id=234.

### Negative Stain sample preparation and data processing

For negative stain, the samples were cross-linked at room temperature for 5 min using 1 mM final concentration of BS3. After cross-linking, 4 μL were applied to a glow discharged continuous carbon grid for 5 minutes, then stained with uranyl formate. A tilted negative stain data set was collected on a Tecnai F20 microscope (FEI) operated at 120 keV and equipped with an Ultrascan 4000 camera (Gatan). Data was collected using Leginon data acquisition software^38^. The CTF parameters were estimated using Gctf (version 1.16) and particles were picked using Gautomatch (version 0.50, from K. Zhang, MRC-LMB, Cambridge) using gaussian blob templates^39^. Data processing was done using Relion (version 3.0)^40^. The negative stain structure (EMD-6834) from Zhang et al. was used as an initial model^41^. Extracted particles were subjected to 2D classification and 3D classification to obtain a homogenous population. Particles that went into the best classes were then refined.

### Cryo-EM sample preparation

For cryo-EM sample preparation we used a Vitrobot Mark IV (FEI). RSC was crosslinked on ice using 1 mM BS3 (Thermo Fisher Scientific) for 15 minutes before 4 μL of sample was applied to either a 2/2 holey carbon or 1.2/1.3 UltrAuFoil grids (Quantifoil) at 4 °C under 100% humidity. The sample was immediately blotted away using Whatman #1 for 2-3 sec at 0 N force and then immediately plunge frozen in liquid ethane cooled by liquid nitrogen. For the RSC-nucleosome complex sample preparation, 4 μL of RSC (2 pmol) and 1 μL of nucleosome (H3K4me3 and H3K(9/14/18)ac) (1 pmol) (Epicypher) were incubated for 10 minutes at 30°C followed by the addition of 0.5 μL of AMPPNP (0.5 nmol) and an additional incubation for 10 minute at 30 °C. The samples were then placed on ice and crosslinked with 1mM BS3 for 15 minutes. RSC-nucleosome samples were prepared in the same way as RSC.

### Cryo-EM data collection

For the RSC sample, frozen grids were clipped and transferred to the autoloader of a Titan Krios electron microscope (Thermo Fischer Scientific) operating at 300 keV (PNCC). Images were recorded with a K3 direct electron detector (Gatan) operating in super-resolution mode at a calibrated magnification of 46,339 (1.079 Å/pixel) and a mean defocus of -1.04 μm with a 0.24 μm standard deviation, using the SerialEM data collection software^42^. 50-frame exposures were taken at 0.06 s per frame, using a dose rate of 11.455 e^-^/pixel/s (0.6 e-Å^2^ per frame), corresponding to a total dose of 40 e^-^Å^-2^ per micrograph (Fig. SX. A total of 8122 movies were collected from a total of 3 grids.

For the RSC-nucleosome sample, frozen grids were clipped and transferred to the autoloader of a Talos Arctica electron microscope (Thermo Fischer Scientific) operating at 200 keV acceleration voltage (UCB). Images were recorded with a K3 direct electron detector (Gatan) operating in super-resolution mode at a calibrated magnification of 43,859 (1.14 Å/pixel) and a mean defocus of -1.66 μm with a 0.41 μm standard deviation, using the SerialEM data collection software^42^. 50-frame exposures were taken at 0.065s per frame, using a dose rate of 11.838 e^-^ /pixel/s (1 e^-^ Å^2^ per frame), corresponding to a total dose of 50 e^-^Å^-2^ per micrograph (Fig. S10). A total of 9190 movie were collected from a single grid.

### Cryo-EM data processing

All data processing was performed using Relion3 (version 3.0)^40^. For the RSC dataset, whole movie frames were aligned and binned by 2 (1.079 Å/pixel) with MotionCor2 to correct for specimen motion The CTF parameters were estimated using Gctf^39,43^. 1,245,498 particles were picked with LoG picker. Particles were extracted binned by 4 (4.316 Å/pixel) and subjected to two- and three-dimensional classification to remove ice and empty picks, which resulted in 1,074,750 particles. The negative stain reconstruction was used as an initial model for 3D classification. The particles were then centered and reextracted bin 1.33 (1.4386 Å/pixel) and refined. The refinement was performed without a mask and resulted in a reconstruction were only the core of the complex was well resolved. The refinement was then continued with a mask around the core. The masked refinement resulted in a 3.8 Å-resolution map of the core. Masked local search 3D classification was performed to select for the best particles. The best class, containing 252,918 particles, was selected and refined, resulting in a 3.4 Å map. Three iterations of CTF refinement, particle polishing and 3D refinement were performed, which resulted in reconstructions at 3.26, 3.21 and 3.18 Å^44^. Local search 3D classification was performed to select the best particles. The best class, containing 192,066 particle images, was selected and further refined, resulting in a 3.14 Å-resolution map. One last iteration of CTF refinement, particle polishing and refinement was performed and led to a reconstructions at 3.07 Å resolution. The core was then subjected to multibody refinement by masking each of the three lobes separately^45^. The arm, head and body refined to 3.23, 3.14, and 2.96 Å respectively. 3D classification of the partially signal subtracted particles was then performed. For the arm lobe a single good class was identified, which when refined resulted in a 3.16 Å map. For the head and body lobes two good classes were found, with one containing several extra helices. The classes containing the extra density were selected and refined. The head lobe refined to 3.07 Å, and the body refined to 3.48 Å. To characterize the two flexible lobes of the complex 3D classification was performed for the 192,066 subset of particles. The classification with four classes resulted in a continuum of states for the leg lobe. The complete tail lobe was only present in two of the four classes, with the tail in two different conformations.

### Model building and refinement

The model for the RSC core was generated by manually building a poly-alanine trace through the final global refinement and multibody maps in COOT^46^. Each of the chains was then identified with the help of blobMapper.py (GitHub: https://github.com/Stefan-Zukin/blobMapper). The resulting coordinate model was iteratively refined using the real space refinement algorithm implemented in PHENIX^47^. Ramachandran, secondary structure, C_β_, and rotamer restraints, as well as bond length and bond angle restraints for the Zn^2+^ ion in the Rsc8 ZZ domain, were used throughout to ensure good model geometry. The final round of refinement comprised 5 rounds of global minimization as well as b-factor refinement and used a resolution limit of 3.1 Angstroms, according to the average resolution of cryo-EM maps used (Extended Data Figure S4e) in order to avoid overfitting of the model. The refinement weight was automatically determined by PHENIX, which monitors the bond length and bond angle deviations to maintain good model geometry and avoids over-fitting of the model to the map^48^. The model was validated using MTRIAGE and MOLPROBITY within PHENIX^48^. The refinement statistics are given in Extended Data Figure S4g and show values typical for structures in this resolution range (MOLPROBITY score = 2.1)^47^. The FSC curve between the model and the map shows good correlation up to 3.2 Angstroms resolution according to the FSC = 0.5 criterion^48^.

The model of the RSC-NCP complex was generated by docking our model of the RSC core, the crystal structure of the Arp module (PDB 4I6M: Arp7, Arp9, Rtt102 and Snf2-HSA) and cryo-EM structure of Snf2-MMTV nucleosome complex bound at the SHL2 with ADP (PDB 6IY2: Snf2, nucleosome)^18,21^. The sequence for the Snf2-HSA helix in 4I6M and Snf2 helicase domain in 6IY2 were aligned to the Sth1 sequence and mutated in COOT^46^.

### Creation of figures and movies

Depiction of molecular models were generated using PyMOL (The PyMOL Molecular Graphics System, version 1.8, Schrödinger), and UCSF ChimeraX developed by the Resource for Biocomputing, Visualization, and Informatics at the University of California, San Francisco, with support from National Institutes of Health R01-GM129325 and the Office of Cyber Infrastructure and Computational Biology, National Institute of Allergy and Infectious Diseases^49^.

### Data availability

The cryo-EM maps and coordinate models have been deposited in the Electron Microscopy Data Bank with the accession codes EMD-XXXX (RSC core), EMD-XXXX (head lobe multibody), EMD-XXXX (body lobe multibody), EMD-XXXX (arm lobe multibody), EMD-XXXX (head lobe classified), EMD-XXXX (body lobe classified), EMD-XXXX (arm lobe classified), EMD-XXXX (RSC-NCP locked) and EMD-XXXX (RSC-NCP swiveled) and in the Protein Data Bank with the accession codes PDB-XXXX (RSC core) and PDB-XXXX (RSC-NCP).

