## Extended_Data_Figures for "Architecture of the chromatin remodeler RSC and insights into its nucleosome engagement"

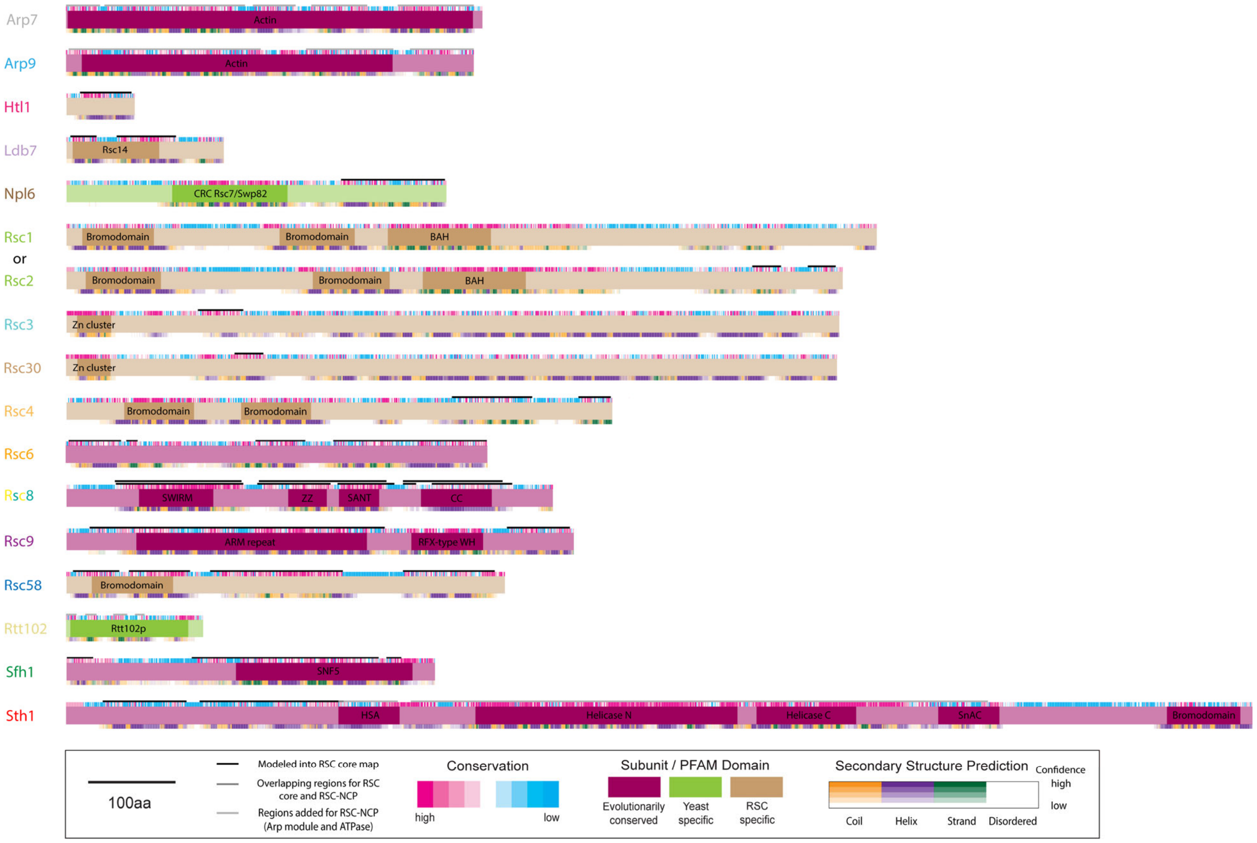


**Fig. S1 | Conservation of RSC proteins with labeled domains and secondary structure predictions.** Regions modeled within our structure are denoted by a black line above the protein. Conservation scores ranging from high to low (pink to blue) were generated using the Consurf server[28,29]. PFAM predicted domains are labeled in bold, with predicted secondary structure based on PSIPRED and DISOPRED3 (purple for helix, green for stand, orange for coil and white for disordered)[30–34].

a

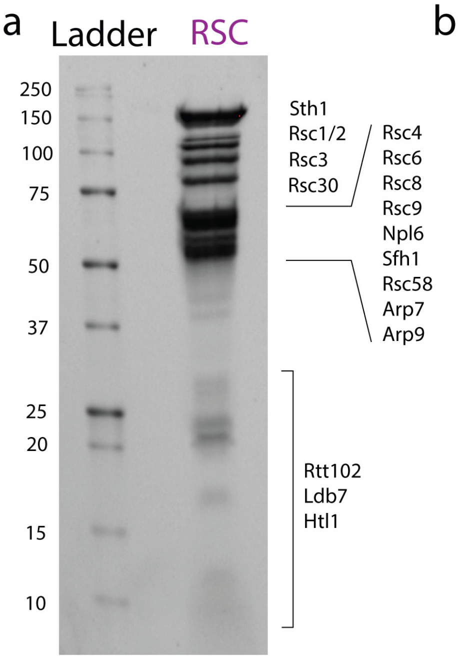


b

| Accession | Description | Peptides | Coverage | Molecular Weight (kDa) |
|---|---|---|---|---|
| P32597 | Sth1 | 50 | 43.9 | 157 |
| P43609 | Rsc8 | 34 | 58.9 | 63 |
| Q06488 | Rsc2 | 30 | 40.3 | 102 |
| Q05123 | Arp9 | 27 | 52.9 | 53 |
| P32832 | Npl6 | 26 | 63.4 | 50 |
| Q03124 | Rsc9 | 24 | 42.9 | 65 |
| Q12406 | Arp7 | 24 | 51.2 | 54 |
| Q07979 | Rsc58 | 22 | 61.8 | 58 |
| P53236 | Rsc1 | 17 | 25.3 | 107 |
| Q06639 | Rsc3 | 16 | 25.3 | 102 |
| P38781; P38780 | Rsc30 | 15 | 25.4 | 101 |
| Q02206 | Rsc4 | 15 | 30.4 | 72 |
| P25632 | Rsc6 | 14 | 36.00 | 54 |
| Q06168 | Sfh1 | 13 | 41.5 | 49 |
| P53330 | Rtt102 | 7 | 65.0 | 18 |
| P38210 | Ldb7 | 5 | 56.1 | 20 |
| Q9URQ5 | Htl1 | 5 | 44.9 | 9 |
| P02407; P14127 | S17-A | 5 | | |
| P10591; P10592 | SSA1 | 1 | | |
| Q05867 | YLR283W | 1 | | |
| P0CX37; P0CX38 | S6-A | 1 | | |
| Q01855 | S15 | 1 | | |
| P11484; P40150 | SSB1 | 1 | | |

c

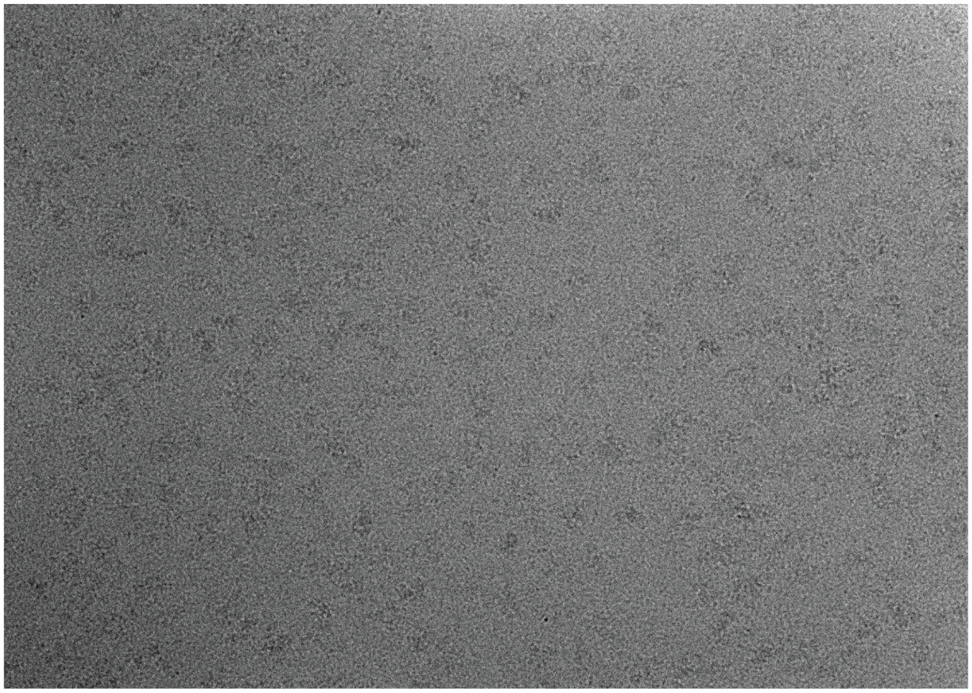

d

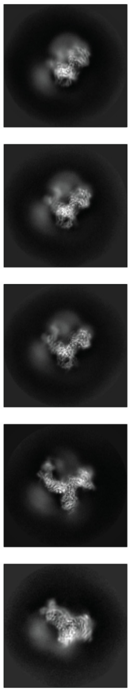

Fig. S2 | Biochemical Characterization of RSC. a, SDS-PAGE of purified S. cerevisiae RSC. b, Mass spectrometry analysis showing the presence of all RSC subunits in our sample. c, Representative cryo-EM micrograph of RSC using open hole grids. d, Exemplar 2D class averages of apo RSC.

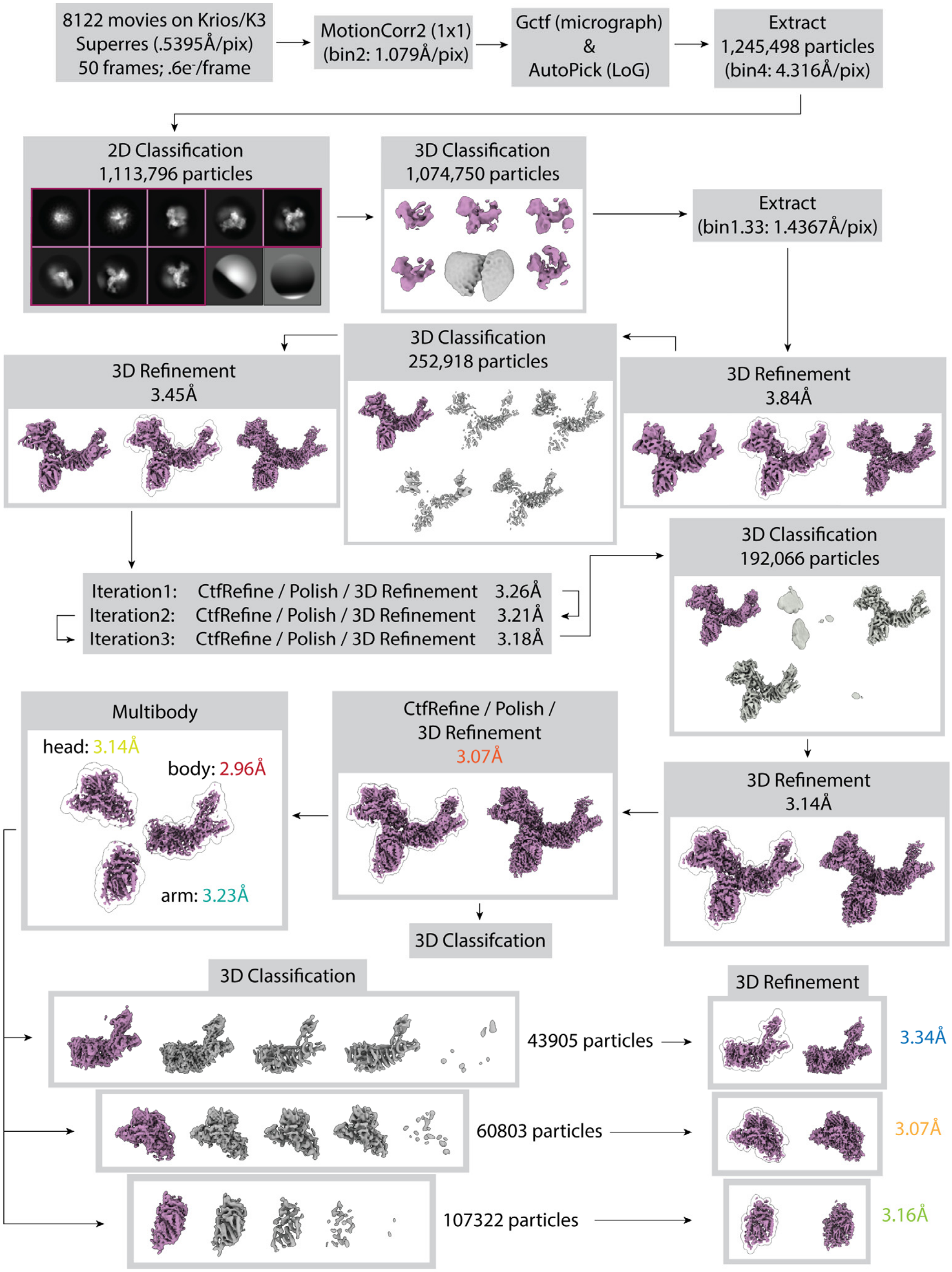


Fig. S3 | Cryo-EM data collection and processing for apo RSC. This processing yielded a map with an overall resolution of 3.07Å for the core. To improve map quality at the periphery of the core, multibody refinement was performed for the head, arm, and body lobes, followed by 3D classification of the partially signal-subtracted particles. For the arm lobe, a single good class was identified, which was refined to a resolution of 3.16 Å. For the head and body lobes, two distinct good classes were found, with one containing several extra helices. The classes containing the extra density were selected and refined. The head lobe refined to 3.07 Å, and the body refined to 3.48 Å.

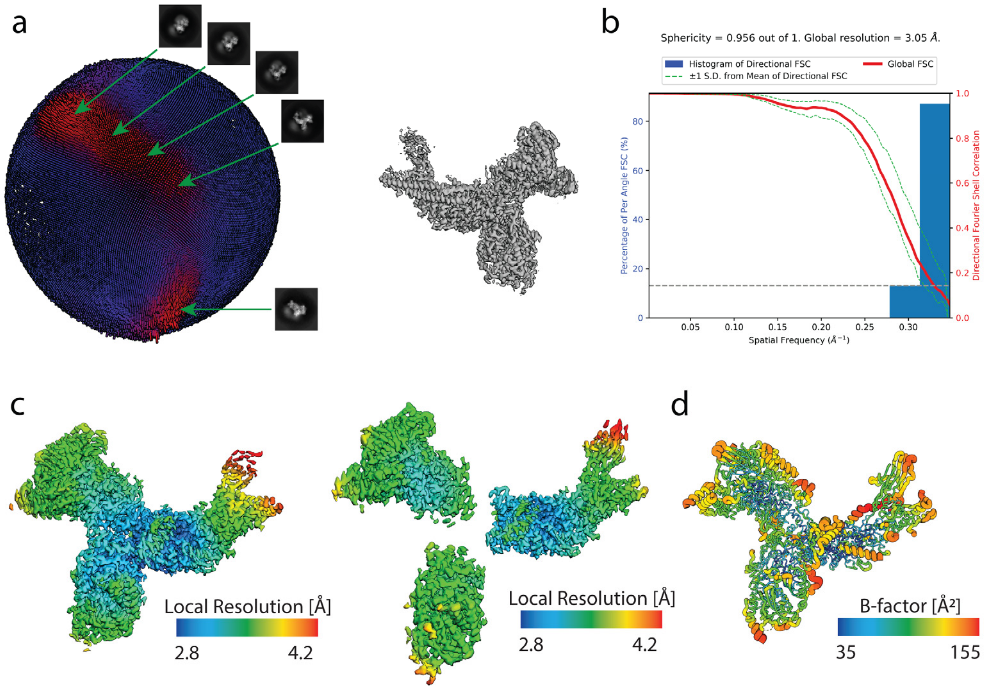


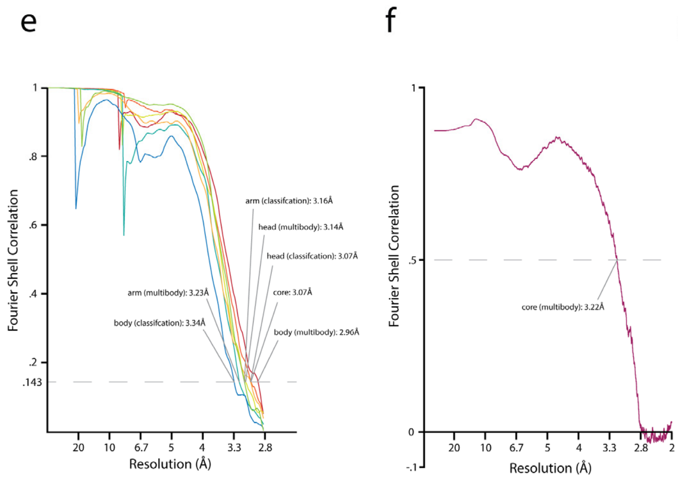


g

| Data Collection, Map and Model Refinement, Validation | | |
|---|---|---|
| **Data Collection** | | |
| Dataset | Apo RSC | RSC-NCP |
| Microscope | Titan Krios | Talos Arctica |
| Stage type | Autoloader | Autoloader |
| Voltage (kV) | 300 | 200 |
| Detector (Mode) | Gatan K3 (super-resolution) | Gatan K3 (super-resolution) |
| Pixel size (Å) | 1.079 (.5395) | 1.14 (.57) |
| Defocus range [μm (mean/STD)] | 2.73 (.91) | 2.80 (.84) |
| Electron dose ($e^-$/Å$^2$) | 40 | 50 |
| **Reconstruction** | | |
| Software | RELION | |
| Particles | 192,066 | |
| Box size (pixels) | 288 | |
| Accuracy rotations (°) | .782 | |
| Accuracy translation (pixels) | .263 | |
| Map resolution (Å) | 3.07 | |
| Map sharpening b-factor (Å$^2$) | -50 | |
| **Coordinate refinement** | | |
| Software | PHENIX | |
| Resolution cutoff (Å) | 3.21 | |
| **Model** | | |
| Residues | | |
| Protein | 3157 | |
| B-factors | | |
| Protein | 76.6 | |
| RMS deviations | | |
| Bond lengths (Å) | .004 | |
| Bond angles (°) | .511 | |
| **Validation** | | |
| Molprobity Score | 6.53 ($89^{th}$) | |
| Molprobity clashscore | 2.11 ($99^{th}$) | |
| Rotamer outliers (%) | 2.45 | |
| Cβ deviations (%) | 0 | |
| Ramachandran plot | | |
| Favored (%) | 92.82 | |
| Allowed (%) | 7.08 | |
| Outliers (%) | .1 | |

Fig. S4 | **a,** Euler angle distribution (left) for the core of RSC (right). Major orientation peaks are marked with corresponding 2D class averages. **b,** 3DFSC plot for apo RSC showing isotropic resolution for the core of RSC[50]. **c,** Maps for the masked refinement and multibody refinement of the RSC core colored in a rainbow spectrum to represent the local resolution (local resolution calculated using Relion)[40]. **d,** Model of the core of RSC, colored according to B-Factor. **e,** FSC plots for the reconstruction in Fig. S3. Resolutions are estimated according to the FSC = .143 criterion (fully independent reconstructions, gold standard). **f,** Model versus map FSC curves for the refined RSC core coordinate model against the summed multibody reconstructions. The FSC = 0.5 criterion indicates a resolution of 3.2 Å for the core of RSC. **g,** Refinement table summarizing data collection settings, reconstruction parameters, and model refinement and validation statistics.

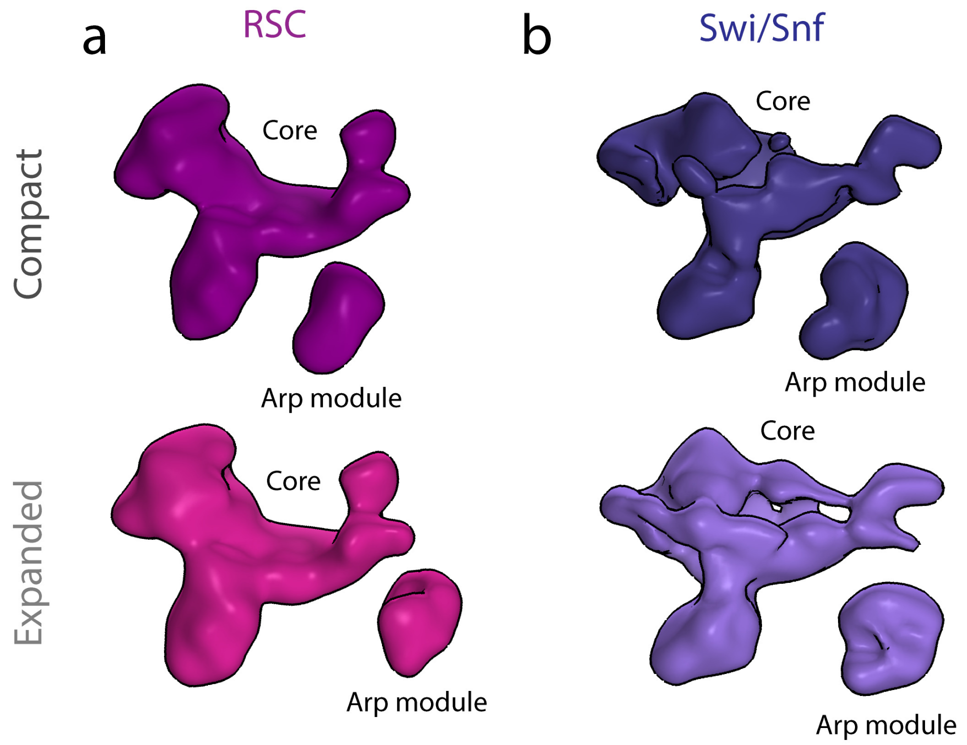


c

| Species | YEAST | | HUMAN | | |
|---|---|---|---|---|---|
| **Complex** | **RSC** | **SWISNF** | **BAF** | **PBAF** | **ncBAF** |
| Conserved | Arp7 | | ACTB | | |
| | Arp9 | | ACTA2 | | |
| | Rsc6 | Swp73 | SMARCD1/2/3 | | |
| | Rsc8 | Swi3 | SMARCC1/2 | | |
| | Sfh1 | Snf5 | SMARCB1 | | |
| | Sth1 | Snf2 | SMARCA2/4 | | |
| | Rsc9 | Swi1 | ARID1A/B | ARID2 | GLTSCR1 |
| Metazoan Specific | | | SMARCE1 | | |
| Yeast Specific | Npl6 | Swp82 | | | |
| | Rtt102 | | | | |
| Complex Specific | Htl1, Ldb7, Rsc1/2, Rsc3, Rsc30, Rsc4, Rsc58 | Snf6, Snf11, Taf14 | DPF2 | PHF10, BRD7, PBRM1 | BRD9 |

Fig. S5 | Negative stain analysis of SWI/SNF and RSC. a, 3D reconstruction of RSC and b, SWI/SNF, both showing a similar three lobed core and a flexible Arp module. c, Homology table for SWI/SNF complexes found in yeast and humans.

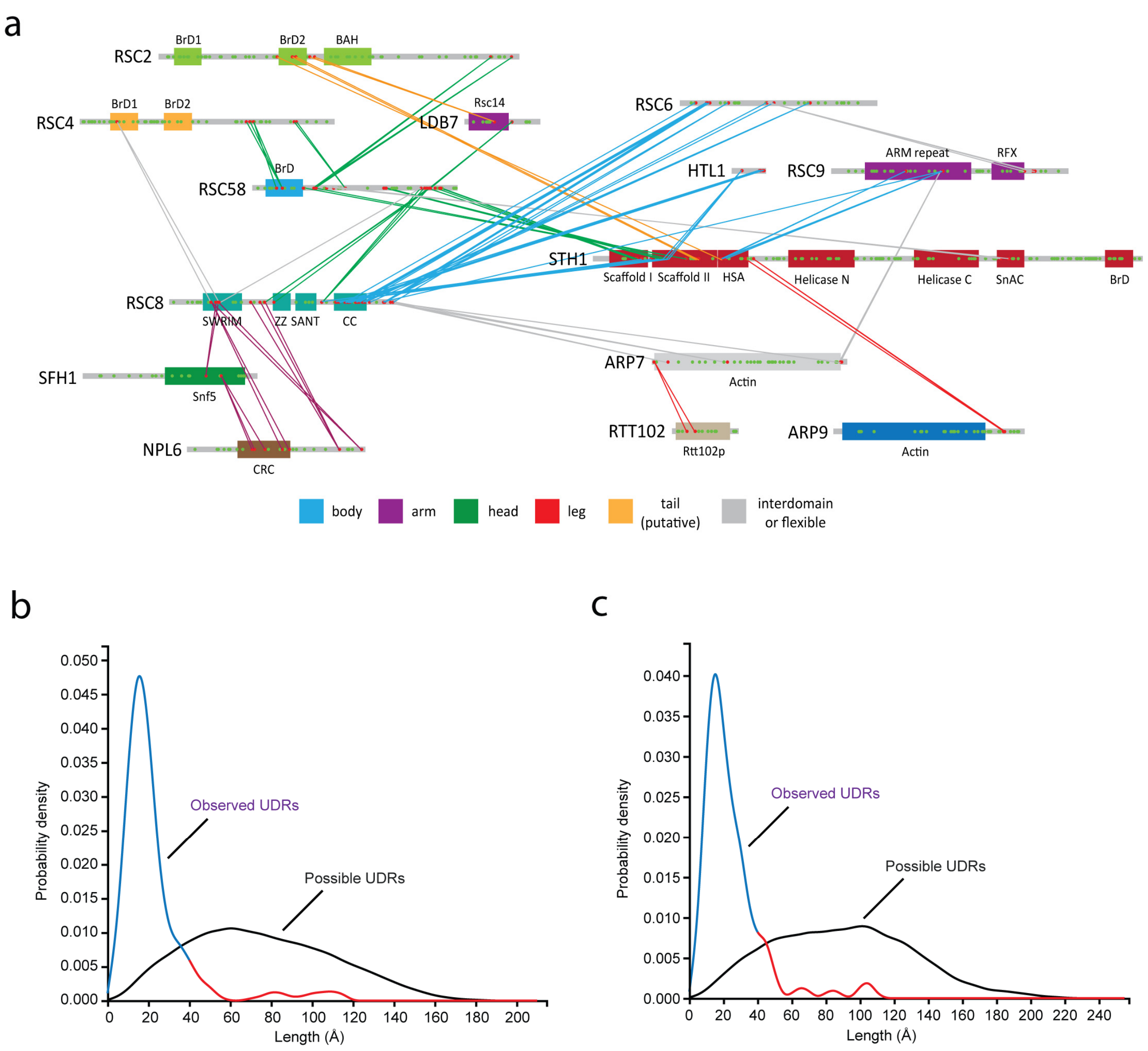


**Fig. S6 | Chemical Crosslinking Mass Spectrometry Analysis of RSC. a**, Protein-protein crosslinks for the RSC complex, plotted on the domain architecture diagram. Only crosslinks with two or more independent crosslinked peptides within 30 amino acids are shown to simplify the display. Crosslinks are colored based on the lobe that they are within. **b,** Probability distribution for the observed and theoretical unique distance restraints (UDRs) for the core of RSC. The observed probability distribution is colored blue (≤38A, possible based on the linker length on BS3) and red (>38A, not possible based on the linker length on BS3). Of the 168 crosslinks that could be mapped onto the structure 151 (90%) were within 38Å. **c**, Same as **(b)** but for the RSC core with the Arp module (Arp7, Arp9, Rtt102, and Sth1-HSA). Of the 233 crosslinks that could be mapped onto the structure 206 (88%) were within 38Å.

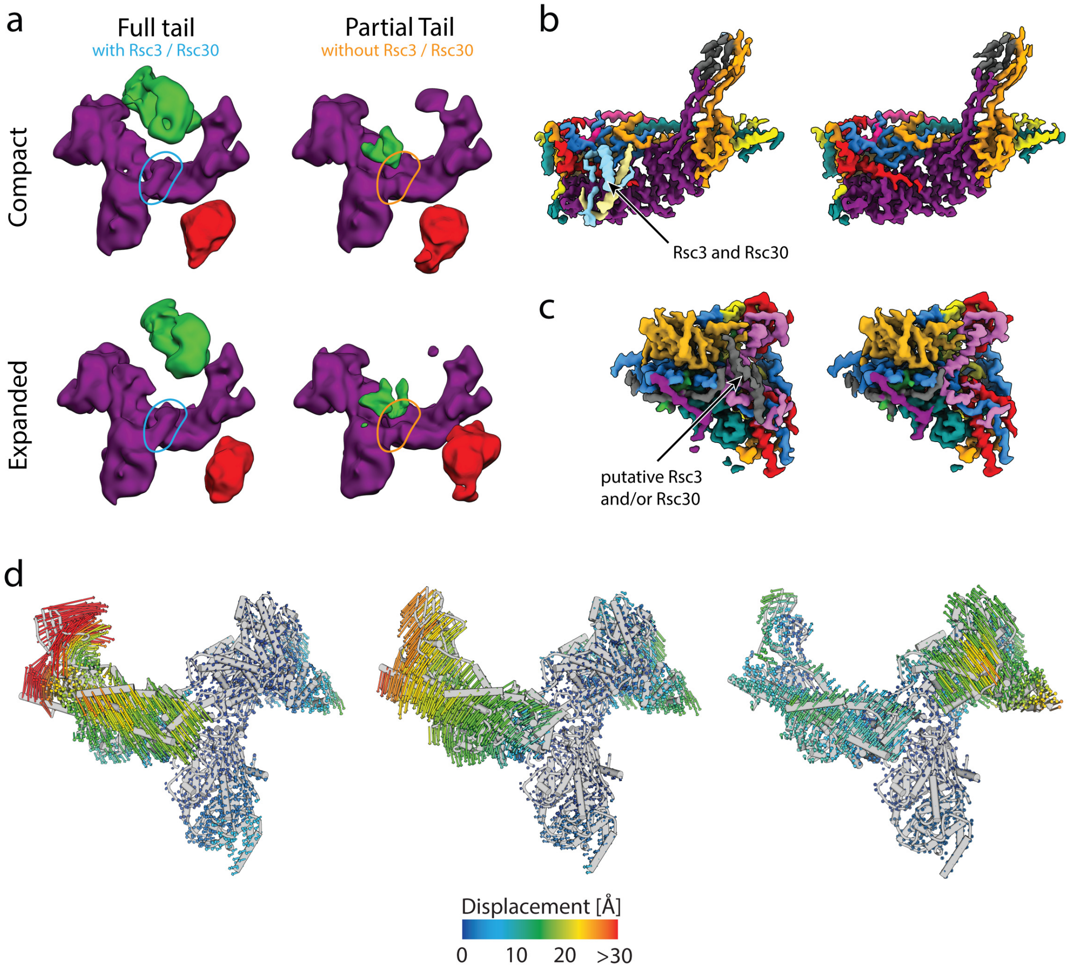


**Fig. S7 | Rsc3 and Rsc30 occupancy in core domain correlates with the presence of the flexible tail.** **a,** Four classes generated from 3D classification using the final high resolution set of particles with no mask and a reference that was low pass filtered to 20Å at every iteration. These classes reveal the flexibility of the tail and leg lobes of RSC and correlate the presence of the tail lobe with Rsc3/Rsc30 occupancy in the core. **b,** Close up of two maps generated from 3D classification where RSC3/30 was either present or absent from the main body of the core. Maps were generated using partial signal subtraction and 3D classification. **c,** Close up of the head lobe of the core from the same maps as **b**, showing the presence or absence of three putative Rsc3/Rsc30 helices. **d,** Cα displacement vectors show the motion of the first three principal components of the multibody refinement. The consensus RSC model was flexibly fit into the first and last volumes of the eigenvalue binned structure and Cα displacement vectors were measured from those resulting structures[51].

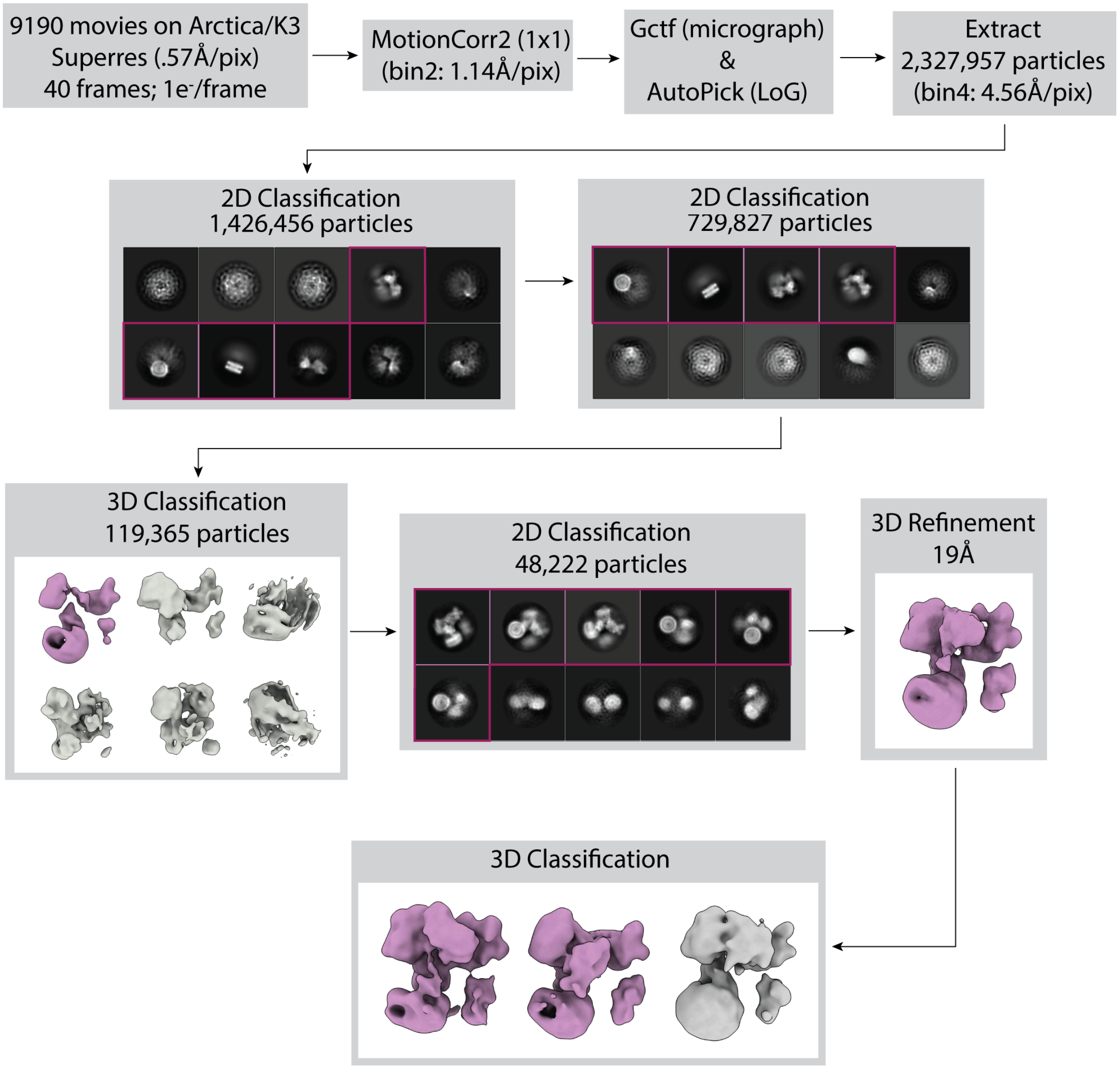


**Fig. S8 | Cryo-EM data collection and processing for RSC-NCP.** This processing scheme yielded an overall reconstruction of RSC bound to a NCP at 19Å. Further classification showed that the tail region shifts towards the NCP in a subset of particles.
